# Susceptibility to gingipains and transcriptomic response to *P. gingivalis* highlights the ribosome, hypothalamus, and cholinergic neurons

**DOI:** 10.1101/2020.08.09.243402

**Authors:** Sejal Patel, Derek Howard, Leon French

## Abstract

*Porphyromonas gingivalis*, a keystone species in the development of periodontal disease, is a suspected cause of Alzheimer’s disease. This bacterium is reliant on gingipain proteases, which cleave host proteins after arginine and lysine residues. To characterize gingipain susceptibility, we performed enrichment analyses of arginine and lysine proportion proteome-wide. Proteins in the SRP-dependent cotranslational protein targeting to membrane pathway were enriched for these residues and previously associated with periodontal and Alzheimer’s disease. These ribosomal genes are up-regulated in prefrontal cortex samples with detected *P. gingivalis* sequences. Other differentially expressed genes have been previously associated with dementia (*ITM2B, MAPI, ZNF267*, and *DHX37*). For an anatomical perspective, we characterized the expression of the *P. gingivalis* associated genes in the mouse and human brain. This analysis highlighted the hypothalamus, cholinergic neurons, and the basal forebrain. Our results suggest markers of neural *P. gingivalis* infection and link the gingipain and cholinergic hypotheses of Alzheimer’s disease.

## Introduction

*Porphyromonas gingivalis* (*P. gingivalis*), a keystone species in the development of periodontal disease is believed to play a pathogenic role in several systemic inflammatory diseases (Fiorillo et al. 2019). This Gram-negative anaerobe is unique in its ability to secrete gingipain proteases, which are its primary virulence factors and are required for its survival *in vivo (Guo, Nguyen, and Potempa 2010). P. gingivalis* is asaccharolytic and uses gingipains to degrade host peptides for nutrition and energy. These gingipain peptidases are cysteine proteases that cleave bonds after arginine (RgpA and RgpB) and lysine (Kgp) (Diego et al. 2014; Bostanci and Belibasakis 2012). These two positively charged amino acid residues facilitate binding of negatively charged nucleic acids (Blanco et al. 2018; Baker et al. 2001), suggesting gingipains will severely disrupt host cell protein-RNA and protein-DNA interactions.

Positive associations between periodontal disease and orodigestive cancer (Olsen and Yilmaz 2019), rheumatoid arthritis (Farquharson, Butcher, and Culshaw 2012; Bingham and Moni 2013), heart disease (Bui et al. 2019), male infertility (Kellesarian et al. 2018), and Alzheimer’s disease have been reported (Singhrao and Olsen 2019; Nadim et al. 2020). Several studies have found links that implicate *P. gingivalis* in these associations. For example, serum *P. gingivalis* antibody levels are a risk factor for orodigestive cancer death (Ahn, Segers, and Hayes 2012), and are more common in rheumatoid arthritis subjects (Mikuls et al. 2009). More directly, *P. gingivalis* expresses enzymes that convert arginine residues to citrulline, which is thought to trigger inflammatory responses in rheumatoid arthritis (Klareskog et al. 2008). In the context of heart disease, *P. gingivalis* DNA has been found in human atherosclerotic plaques (Nakano et al. 2006; Haraszthy et al. 2000) and gingipains modify high- and low-density lipoproteins (Lönn et al. 2018). More specifically, gingipains are able to fragment Apolipoprotein E. This fragmentation by Arg-specific gingipains occurs at arginine residues in the protein (Lönn et al. 2018). *APOE* is the strongest genetic riskfactorfor late-onset Alzheimer’s disease, differences between the genetic variants increase the number of arginine residues at two positions. The lowest risk APOE2 isoform encodes only cysteines at these positions, while APOE3 contains one arginine and APOE4, which confers the highest risk contains two (C.-C. Liu et al. 2013). It has been proposed that this variable susceptibility to gingipain cleavage of APOE results in differing amounts of neurotoxic fragments (Dominy et al. 2019). This connection to genetic risk is supported by significant evidence of *P. gingivalis* and specifically gingipains in Alzheimer’s disease pathogenesis (Dominy et al. 2019; Ilievski et al. 2018; Poole et al. 2015; Ishida et al. 2017; Singhrao and Olsen 2019). This evidence linking *P. gingivalis* to several chronic diseases combined with the direct link between arginine count and genetic risk for Alzheimer’s disease motivated our genome- and brain-wide study of gingipain susceptibility.

In this study, we sought to determine which human proteins are susceptible to gingipain cleavage by characterizing proteins with high proportions of arginine and lysine. Motivated by evidence linking *P. gingivalis* to Alzheimer’s disease, we tested if the genes encoding these proteins are differentially expressed in brain tissue with detected *P. gingivalis* RNA. We then performed a neuroanatomical analysis to determine which brain regions are enriched for expression of genes associated with *P. gingivalis* detection and have high arginine and lysine proportion to better understand tissue specific susceptibility. We extend these analyses to a single-cell atlas of the mouse nervous system to identify cell-types that may be specifically susceptible to gingipains.

## Methods

### Amino acid distribution analysis

Translated human protein sequences were obtained from GENCODE version 32 (Frankish et al. 2019). Amino acid proportions were mean averaged across multiple transcripts that were annotated to the same gene symbol. Protein sequences annotated to more than one gene were removed to prevent amplification of single sequences in the following enrichment analyses.

### Gene Ontology enrichment analysis

The Gene Ontology (GO) provides gene-level annotations that span specific cellular components, biological processes, and molecular functions (Ashburner et al. 2000). These annotations, defined by GO terms, were required to have annotations for 10 to 200 tested genes (6,807 unique GO groups annotating 14,655 unique genes). To test for enrichment, we sorted the genes from the most enriched to the most depleted proportions of arginine and lysine residues. Within this ranking, the area under the receiver operating characteristic curve (AUC) was used to test for gene ontology terms that are enriched in either direction (overexpressed: AUC > 0.5, underexpressed: AUC < 0.5). The Mann–Whitney U test was used to determine statistical significance with FDR correction for the GO groups tested. We used GO annotations from the GO.db and org.Hs.eg.db packages in R, version 3.8.2, which were dated April 24, 2019 (Carlson 2016a, [b] 2016).

### Prefrontal cortex differential expression

Gene expression profiles from a postmortem study of Parkinson’s disease were used to test for gene expression differences in tissue samples with *P. gingivalis* reads (Dumitriu et al. 2016). To exclude confounding effects of disease processes, we limited our analyses to the 60 neurologically normal control samples of this study. As detailed by Dumitriu et al., prefrontal cortex samples were profiled with Illumina’s HiSeq 2000 sequencers. All samples were from males of European ancestry. Neuropathology reports were used by Dumitriu et al. to exclude brains with Alzheimer’s disease pathology beyond that observed in normal aging.

We used the Sequence Read Archive analysis tool to identify samples with sequencing reads that align to the *P. gingivalis* genome (Leinonen et al. 2011).To test for differential expression, we obtained the normalized count matrix for GSE68719 (GSE68719_mlpd_PCG_DESeq2_norm_counts.txt.gz). To identify genes differentially expressed in samples with detected *P. gingivalis*, we modelled log(1+expression) with an ordinary least squares linear model with covariates for age, RIN, PMI, and total bases read. Gene ontology enrichment analyses was performed using the same AUC method described above.

To examine differential expression of cell-type markers, we used the top marker genes obtained from a single cell study of the adult human temporal cortex (Darmanis et al. 2015). This study used gene expression profiles to cluster cells into astrocyte, neuron, oligodendrocyte, oligodendrocyte precursor, microglia and endothelial groups. These marker genes were used to calculate AUC values.

### Gene expression processing for anatomical enrichment analysis

The Allen Human Brain Atlas was used to determine regional enrichment of the genes identified in the differential expression analyses throughout the brain (Hawrylycz et al. 2012). Thus, in contrast to the differential analyses that focused on samples from the cerebral cortex, this approach used the whole brain, and expression profiles were summarized to named brain regions. For each donor, samples mapping to the same-named brain region were mean-averaged to create a single expression profile for each region. Values from analogous named brain regions of both hemispheres were pooled because no significant differences in molecular architecture have been detected between the left and right hemispheres (Miller et al. 2014). Expression levels of the 48,170 probes were then summarized by mean averaging for each of 20,778 gene transcripts. Expression values were next converted to ranks within a named brain region, and then z-score normalized across brain regions. For the analyses that combine across donors, z-scores for each donor were averaged to obtain a single gene-by-region reference matrix of expression values.

### Mouse nervous system gene expression data

The RNA sequencing data that were used to characterize gene expression across the mouse nervous system were obtained from Zeisel et al. (2018). This dataset of more than 500,000 isolated cells from 19 defined regions was obtained from male and female mice that ranged in age from 12 to 56 days old. Following quality control, Zeisel et al. identified 265 clusters of cells. These clusters are thought to represent different cell types. Aggregate expression values for the identified transcriptomic cell-type clusters were obtained from the Mousebrain.org downloads portal. These values were log(expression+1) transformed and then z-scored at the gene level. We restricted our analysis to genes with human homologs by filtering mouse gene symbols for those that have human homologs (O’Leary et al. 2016). Expression z-scores are then ranked genome-wide within a defined cell-type cluster.

### Neuroanatomical and cell-type cluster enrichment analysis

To calculate enrichment for gene sets of interest within a brain region or a cell type, z-scores in the processed expression matrices are first ranked. This provides a genome-wide list for each cell type or brain region with high ranks marking specific and low for depleted expression. We project our genes of interest into these ranked lists to determine enrichment. The AUC statistic was used to quantify if the genes of interest are more specifically expressed (ranked higher) in this sorted list of genes for a specific region or cell cluster. In this context, AUC > 0.5 means the associated genes have specific or enriched expression in a brain region/cell-type cluster. For AUC < 0.5, the reverse is true, with a bias toward lower relative expression. The Mann-Whitney U test was used to test the statistical significance, and the FDR procedure was used to correct for testing of multiple brain regions or cell clusters within a dataset.

#### Availability of data and code

Scripts, supplementary tables, and data files for reproducing the analyses are available online at https://figshare.com/articles/dataset/Susceptibility_to_gingipains_and_transcriptomic_response_to_P_gingivalis_highlights_the_ribosome_hypothalamus_and_cholinergic_neurons/12782576 and <Github link forthcoming>.

## Results

### Genome-wide search reveals a range of arginine and lysine proportions

The average proportion of arginine or lysine residues was 11.4% across the 20,024 human protein-coding genes in our analysis. A broad range was observed, with 95% of genes having proportions between 5.5% and 19.4% (full listing in Supplement Table 1). The gene with the highest proportion was Ribosomal Protein L41 (*RPL41*, 68%) and joined three other ribosomal proteins within the top ten list (*RPL39, RPS25*, and *RPL19*). Ranked second and third are genes known to function in sperm DNA compaction (*PRM1* [47.1%] and *TNP1* [38.2%]). This high proportion of arginine and lysine in these top proteins suggests they are highly susceptible to gingipain cleavage.

### Ribosomal and DNA packaging genes are enriched for arginine and lysine residues

To characterize the proteins with high arginine and lysine residues genome-wide, we performed a GO enrichment analysis. Of the 6,806 tested GO groups, 881 are enriched for higher proportions of arginine and lysine after multiple test correction (top ten in Table 1, full listing in Supplement Table 2). In agreement with inspection of the top ten proteins, genes encoding parts of the ribosomal subunit are the most strongly enriched with one in five residues being arginine or lysine on average (AUC = 0.87, p_FDR_ < 10^−58^). The remaining top groups largely contain ribosomal protein genes. Ranked 15th are genes annotated to the nucleosome, which includes many histones and the two above mentioned genes involved in DNA packaging in sperm (60 genes, 20.9% average proportion, AUC = 0.926, p_FDR_ < 10^−29^). The high enrichment for this set is partially due to 9 highly similar protein sequences from a histone microcluster that has the same arginine and lysine proportion (22.2%). Overall, this enrichment for positively charged arginine and lysine residues mirrors their known ability to facilitate the binding of DNA and ribosomal RNA (Baker et al. 2001; Lott, Wang, and Nakazato 2013).

**Table 1.** Top GO groups enriched for high arginine + lysine proportion.

| Title | ID | Gene count | AUC | p | pFDR |
| --- | --- | --- | --- | --- | --- |
| ribosomal subunit | GO:0044391 | 167 | 0.871 | 1.78E-61 | 1.16E-57 |
| structural constituent of ribosome | GO:0003735 | 114 | 0.924 | 3.16E-55 | 1.03E-51 |
| protein targeting to ER | GO:0045047 | 97 | 0.893 | 6.57E-41 | 1.43E-37 |
| establishment of protein localization to endoplasmic reticulum | GO:0072599 | 101 | 0.881 | 4.38E-40 | 7.16E-37 |
| cytosolic ribosome | GO:0022626 | 93 | 0.891 | 5.79E-39 | 6.79E-36 |
| SRP-dependent cotranslational protein targeting to membrane | GO:0006614 | 92 | 0.893 | 6.23E-39 | 6.79E-36 |
| large ribosomal subunit | GO:0015934 | 104 | 0.869 | 7.96E-39 | 7.44E-36 |
| cotranslational protein targeting to membrane | GO:0006613 | 95 | 0.879 | 1.58E-37 | 1.29E-34 |
| nuclear-transcribed mRNA catabolic process, nonsense-mediated decay | GO:0000184 | 116 | 0.842 | 2.54E-37 | 1.84E-34 |
| translational initiation | GO:0006413 | 181 | 0.762 | 3.66E-34 | 2.39E-31 |

Within the top ten most enriched GO groups, ‘SRP-dependent cotranslational protein targeting to membrane’ appears to be the most specific with the lowest number of annotated genes. These genes are primarily components of the cytosolic ribosome that facilitate translation into the endoplasmic reticulum (ER). For brevity, we refer to this GO group as the ‘ER translocation’ genes. This specific GO group has been previously associated with disorders that *P. gingivalis* is believed to play a pathogenic role.

Specifically, a spatial transcriptomics study reported higher expression of the ER translocation genes in inflamed areas of periodontitis-affected gingival connective tissue compared to non-inflamed areas (Lundmark et al. 2018). A second study that examined peri-implant soft tissue found that ER translocation genes are expressed at higher levels in diseased mucosa samples (Ludden 2015). In the context of Alzheimer’s disease, the ER translocation genes are upregulated across neuroinflammation, in cases from a Caribbean Hispanic postmortem study and in regions of Alzheimer’s disease associated hypometabolism (Patel et al. 2020; Felsky et al. 2020). Given this upregulation of the ER translocation genes in periodontal and Alzheimer’s disease, we pursued further characterization of the ER translocation genes.

Genome-wide, the ER translocation genes have a high proportion of arginine and lysine, but it’s not clear if other residues are enriched. To determine the specificity of this enrichment, we tested proportions of other amino acids. For single residues, lysine (AUC = 0.89) and arginine (AUC = 0.76) are the most enriched in the ER translocation genes, followed by valine and isoleucine (AUC = 0.64 and 0.63, respectively) (Supplement Table 3). For pairs, Figure 1 shows the AUC values of the 190 possible amino acid combinations. Only the lysine and valine proportion have a higher enrichment score, which is slightly higher (AUC = 0.896 compared to 0.894 for arginine + lysine). Still, this combination is less frequent (18.3% of residues compared to 20.4%). Overall, the ER translocation genes are strongly and specifically enriched for arginine and lysine residues, suggesting they are particularly susceptible to cleavage by gingipains.

**Figure 1.**
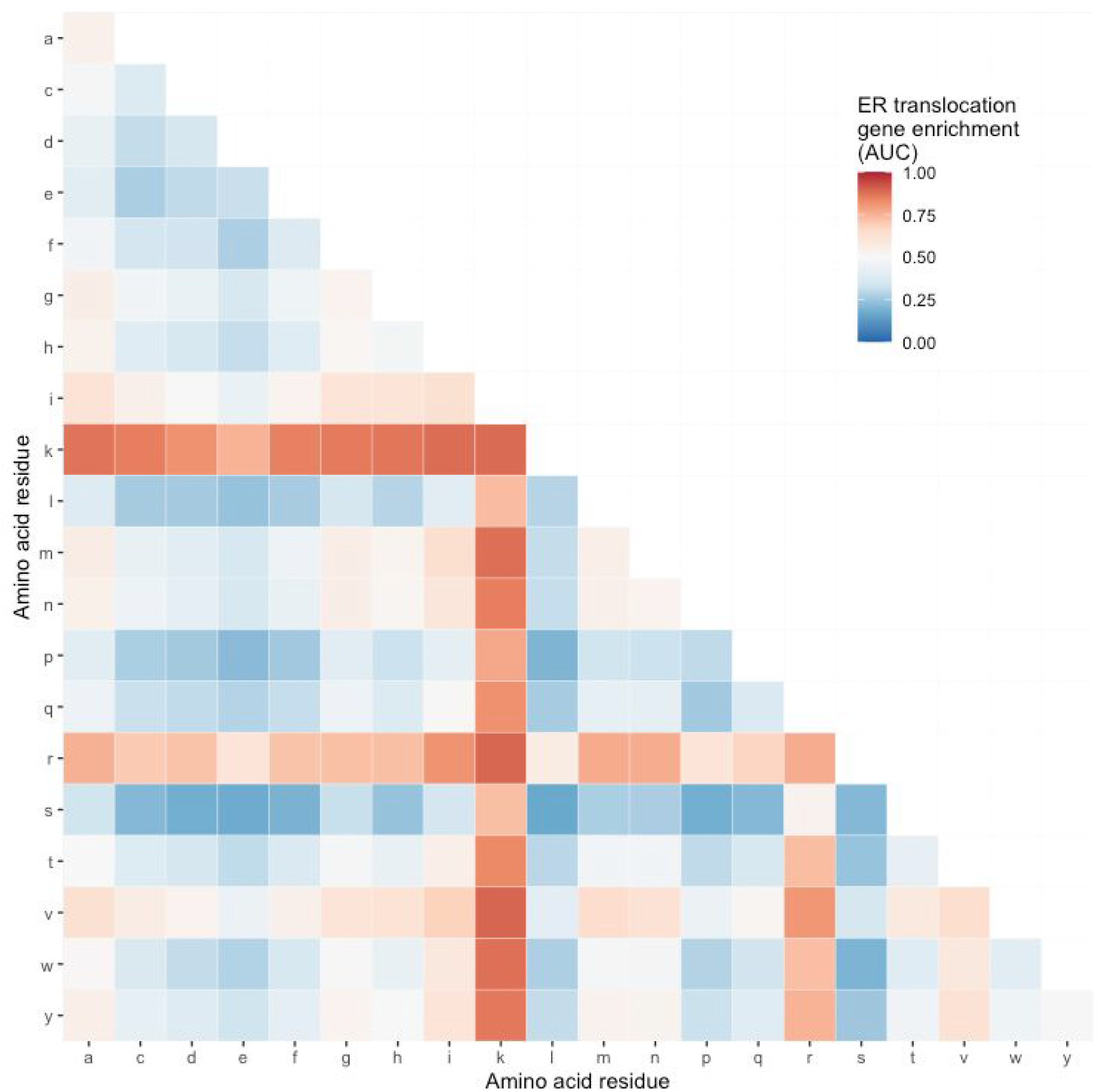
Heatmap of ER translocation gene enrichment for proportions of different amino acid pairs. Relative to all other genes, proportions of specific amino acid pairs for the ER translocation genes range from high (red) to low (blue).

### Hundreds of genes are differentially expressed in brain tissue with *P. gingivalis* reads

Guided by the findings of the upregulation of ER translocation genes in the context of Alzheimer’s disease, we tested for direct associations with *P. gingivalis* in postmortem brain tissue. We examined control samples that lacked any neurodegenerative disease pathology to remove any late-stage signals of Parkinson’s or Alzheimer’s disease. In this all-male dataset, age ranged from 46 to 97 years old. Within the 44 prefrontal cortex samples, *P. gingivalis* sequencing reads were detected in ten. In contrast, reads from the two other bacteria in the red complex, which is associated with severe periodontal disease were not detected (*Tannerella forsythia* and *Treponema denticola*). There was no difference between age, RNA integrity number, and postmortem interval between the samples with and without detected *P. gingivalis* reads (all p > 0.33). In contrast, there was a significant difference in the number of bases sequenced, with more reads in the samples with detected *P. gingivalis*. All four of these variables were covariates in our differential expression model.

In total, 2,189 of the 15,936 tested genes were differentially expressed. More genes were down-than up-regulated in samples with detected *P. gingivalis* reads (1247 versus 942). Arginine and lysine proportion was different between these two sets of genes with an average proportion of 12.8% for the up-regulated genes versus 11.4% for the down-regulated genes (p < 10^−16^). The top ten most up- and down-regulated genes are provided in Table 2 (full listing in Supplement Table 4). The third most up-regulated gene is signal recognition particle 9 (*SRP9*, p_FDR_ < 0.0013, Figure 2A), a member of the ER translocation gene set that forms a heterodimer with the next ranked SRP gene (*SRP14*, p_FDR_ < 0.02) (Birse et al. 1997). The strongest downregulated genes include putative RNA helicases (*DHX30, DHX37*) and other genes involved in the regulation of global gene transcription (*ZNF696*).

**Table 2.** Top ten most up- and down-regulated differentially expressed genes in the prefrontal cortex with detected *P. gingivalis* reads.

| name | symbol | estimate | $P_{\text{FDR}}$ |
| --- | --- | --- | --- |
| integral membrane protein 2B | ITM2B | 0.446 | 0.00107 |
| transmembrane and coiled-coil domains 1 | TMCO1 | 0.357 | 0.00129 |
| signal recognition particle 9 | SRP9 | 0.556 | 0.00129 |
| eukaryotic translation initiation factor 3 subunit M | EIF3M | 0.273 | 0.00129 |
| RAB28, member RAS oncogene family | RAB28 | 0.463 | 0.00133 |
| anti-silencing function 1A histone chaperone | ASF1A | 0.411 | 0.00134 |
| inner mitochondrial membrane peptidase subunit 1 | IMMP1L | 0.555 | 0.00134 |
| MNAT1 component of CDK activating kinase | MNAT1 | 0.389 | 0.0014 |
| zinc finger protein 267 | ZNF267 | 0.754 | 0.00159 |
| mitochondrial calcium uptake 2 | MICU2 | 0.393 | 0.00165 |
| ..... |  |  |  |
| transmembrane protein 63B |  | -0.417 | 0.000998 |
| ubiquitin like modifier activating enzyme 1 | UBA1 | -0.372 | 0.000998 |
| dynein cytoplasmic 1 heavy chain 1 | DYNC1H1 | -0.446 | 0.000998 |
| trafficking protein particle complex 9 | TRAPPC9 | -0.406 | 0.000963 |
| ATP binding cassette subfamily F member 3 | ABCF3 | -0.362 | 0.000963 |
| fatty acid synthase | FASN | -0.701 | 0.000812 |
| transmembrane protein 8B | TMEM8B | -0.4 | 0.000812 |
| DEAH-box helicase 37 | DHX37 | -0.513 | 0.000745 |
| zinc finger protein 696 | ZNF696 | -0.311 | 0.000745 |
| DExH-box helicase 30 | DHX30 | -0.4 | 0.000745 |

**Figure 2.**
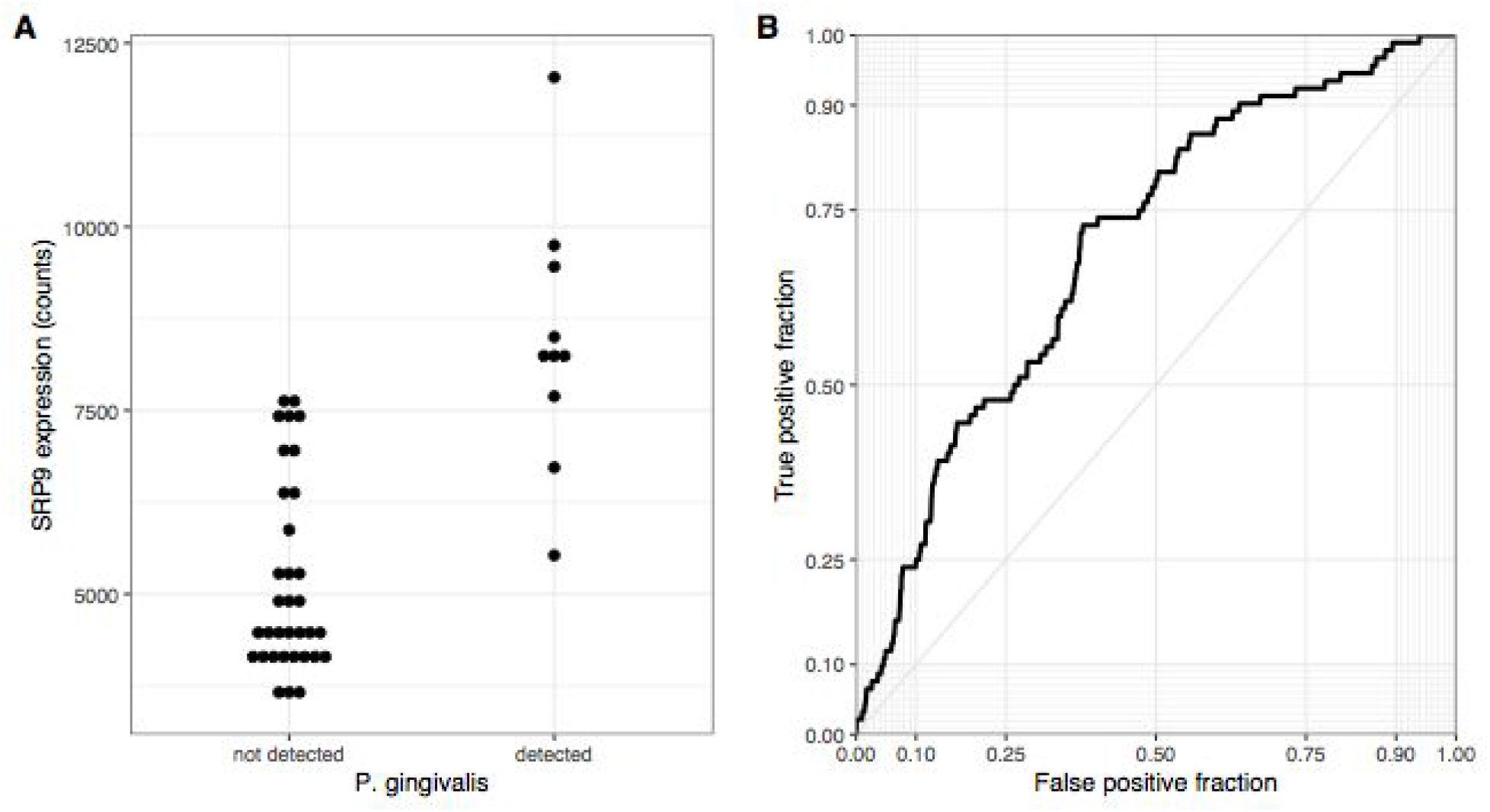
Visualization of differential expression results. A) Dotplot of *SRP9* expression is plotted for samples with and without *P. gingivalis* reads. B) ROC for the ER translocation genes when all genes are ranked according to direction and significance (signed log(p-value)).

### ER translocation genes are enriched for upregulation in samples with *P. gingivalis* reads

To summarize the hundreds of differentially expressed genes, we again used the AUC metric. Genes were ranked from the most up- to down-regulated in the samples with detected *P. gingivalis*. Of the 6,538 tested GO groups, 23 were significantly up-regulated, and 49 were down-regulated after multiple test correction (full listing in Supplement Table 5). Across this ranking, ER translocation genes were strongly up-regulated and is the fourth most significant GO group (AUC = 0.698, p_FDR_ < 10^−7^). Similar GO groups that primarily contain genes encoding ribosomal proteins make up the other top ten up-regulated groups. Of these ten, all are in the top ten list of groups that are enriched for arginine and lysine residues except the ‘protein localization to endoplasmic reticulum’ and ‘protein targeting to membrane’ groups. Within the top ten most down-regulated GO groups, ‘homophilic cell adhesion via plasma membrane adhesion molecules’ ranked first, followed by several synapse associated groups and ‘ATP biosynthetic process’ (all p_FDR_ < 0.005). In the context of amino acid residues, we note that genes annotated to ‘aminoacyl-tRNA ligase activity’ are also down-regulated (AUC = 0.33, p_FDR_ < 0.05). In summary, samples with detected *P. gingivalis* reads have higher expression of ER translocation genes.

Given the down-regulation of synapse genes, we next tested if cell-type specific markers are differentially expressed. Of the six cell-types, only genes marking endothelial cells (AUC = 0.805, p_FDR_ < 0.00005) and astrocytes (AUC = 0.682, p_FDR_ < 0.02) were enriched, with higher expression in samples with *P. gingivalis* reads. In contrast, markers of oligodendrocyte precursors, oligodendrocytes, neurons, and microglia were not enriched. Specifically, the 20 neuronal markers are equally split between up- and down-regulated with no genes reaching statistical significance at an alpha of 0.05 (uncorrected). In summary, differential expression of cell-type markers suggests changes in endothelial and astrocyte cell states or proportions but not neurons.

### Genes up-regulated in samples with *P. gingivalis* reads are highly expressed in the anterior hypothalamic area

To further characterize the genes associated with *P. gingivalis*, we determined which brain regions are enriched for their expression. While these genes were identified in samples from the prefrontal cortex, it is believed neocortical degeneration doesn’t occur until later stages of Alzheimer’s disease (Braak and Braak 1991). Testing for neuroanatomical enrichment may reveal other regions of interest. In this analysis, for a given brain region, the genome was ranked from the most specifically expressed gene to the most depleted gene, relative to the rest of the brain. Similar to the preceding analyses, we use the AUC metric to test if genes of interest rank higher in this list to determine region-specific expression. In total, 82 of 232 tested brain regions are enriched for high expression of the genes up-regulated in samples with detected *P. gingivalis* (Supplement Table 6). The anterior hypothalamic area most specifically expresses these genes (AUC = 0.740, pFDR < 10^−120^). Given the differences in regions sampled across the six brains, we performed this analysis for each donor individually. The anterior hypothalamic area is the 7th ranked brain region for selective expression of genes up-regulated in samples detected with *P. gingivalis* reads in donor 10021/H0351.2002, and is ranked 3rd of 185 in donor 14380/H0351.1012. Within the top ten most enriched regions, two other hypothalamic regions appear (Table 3, AUCs > 0.68, both p_FDR_ < 10^−76^). Notably, most of the top regions border ventricles (medial habenular nucleus, thalamic paraventricular nuclei, substantia innominata, central gray of the pons, paraventricular nucleus of the hypothalamus, and septal nuclei).

**Table 3.** Top ten regions enriched for higher expression of genes up-regulated in samples with detected *P. gingivalis* reads.

| | AUC | p | $p_{FDR}$ |
| --- | --- | --- | --- |
| anterior hypothalamic area | 0.740 | 1.16E-135 | 2.68E-133 |
| medial habenular nucleus | 0.702 | 2.96E-97 | 3.44E-95 |
| paraventricular nucleus of the hypothalamus | 0.684 | 4.60E-81 | 3.56E-79 |
| lateral hypothalamic area, anterior region | 0.683 | 1.85E-79 | 1.08E-77 |
| septal nuclei | 0.682 | 6.28E-79 | 2.92E-77 |
| substantia innominata | 0.679 | 8.62E-77 | 3.33E-75 |
| central gray of the pons | 0.675 | 3.77E-73 | 1.25E-71 |
| midbrain reticular formation | 0.672 | 2.03E-70 | 5.88E-69 |
| paraventricular nuclei, right of thalamus | 0.658 | 1.00E-59 | 2.59E-58 |
| paraventricular nuclei, left of thalamus | 0.656 | 1.26E-58 | 2.92E-57 |

### Cells in the mouse peripheral nervous system and cholinergic neurons highly express genes up-regulated in samples with *P. gingivalis* reads

We next characterized the expression patterns of the genes up-regulated in samples with detected *P. gingivalis* at a finer resolution in the mouse nervous system (Zeisel et al. 2018). Similar to the regional analyses above, we ranked each gene from the most specific to depleted expression for each transcriptomic cell type cluster. The 873 mouse homologs of the up-regulated genes are enriched for specific expression in 79 of the 265 tested transcriptomic cell type clusters (Supplement Table 7). The top two enriched clusters are nitrergic enteric neurons (both AUC > 0.67, p_FDR_ < 10^−65^). The next three most enriched are cholinergic neurons with probable locations listed as the sympathetic ganglion or myenteric plexus of the small intestine (all AUC > 0.66, p_FDR_ < 10^−56^). The top 19 most enriched clusters are located in the peripheral nervous system. Focusing on the central nervous system, the top two clusters described as “Afferent nuclei of cranial nerves VI-XII” and “Cholinergic neurons, septal nucleus, Meissnert and diagonal band” (both AUC < 0.59, p_FDR_ < 10^−19^). Broadly, cells in the mouse peripheral nervous system and cholinergic neurons strongly express genes that are up-regulated in samples with detected *P. gingivalis*.

### ER translocation genes are highly expressed in the substantia innominata

To extend our results beyond protein level susceptibility, we next identified brain regions that express high levels of the ER translocation genes to determine neuroanatomical susceptibility related to *P. gingivalis*. While our preceding analyses used genes identified from brain tissue samples, the ER translocation genes were identified from analyses of protein sequences alone. This provides independence from tissue- and state-specific expression. In a combined analysis of all six brains, 83 of the 232 brain regions showed overexpression of the ER translocation genes. The substantia innominata was top-ranked with the most specific expression of the ER translocation genes (AUC = 0.846, p_FDR_ < 10^−28^, full listing in Supplement Table 8). Of the nine remaining top 10 regions, seven are located near the substantia innominata: internal and external globus pallidus, substantia nigra pars reticulata, the septal nuclei, nucleus accumbens, subcallosal cingulate gyrus, head of the caudate nucleus (AUCs > 0.774, all p_FDR_ < 10^−18^). Of the 34 assayed cerebellar cortex regions, all are significantly enriched for expression of the ER translocation genes. Within specific brains, the substantia innominata is the 10th ranked brain region (of 194) for selective expression of ER translocation associated genes in donor 10021/H0351.2002, and is ranked 3rd of 182 in donor 9861/H0351.2001. There are no samples from the substantia innominata in the other four donors, but its constituent nuclei are. Testing for anatomical enrichment of the ER translocation associated genes in these nuclei reveals a highly heterogeneous pattern (Figure 3). A key characteristic of the substantia innominata is a high proportion of cholinergic neurons (M. M. Mesulam et al. 1983). Alzheimer’s disease has been previously associated with cholinergic neuron loss in the basal forebrain and deficits in choline O-acetyltransferase (encoded by *CHAT*) (Sims et al. 1983; Hampel et al. 2019; DeKosky et al. 1992). Figure 3 shows the relationships between CHAT gene expression and ER translocation genes across the brain, marking the substantia innominata as having high expression of both *CHAT* and the ER translocation genes.

**Figure 3.**
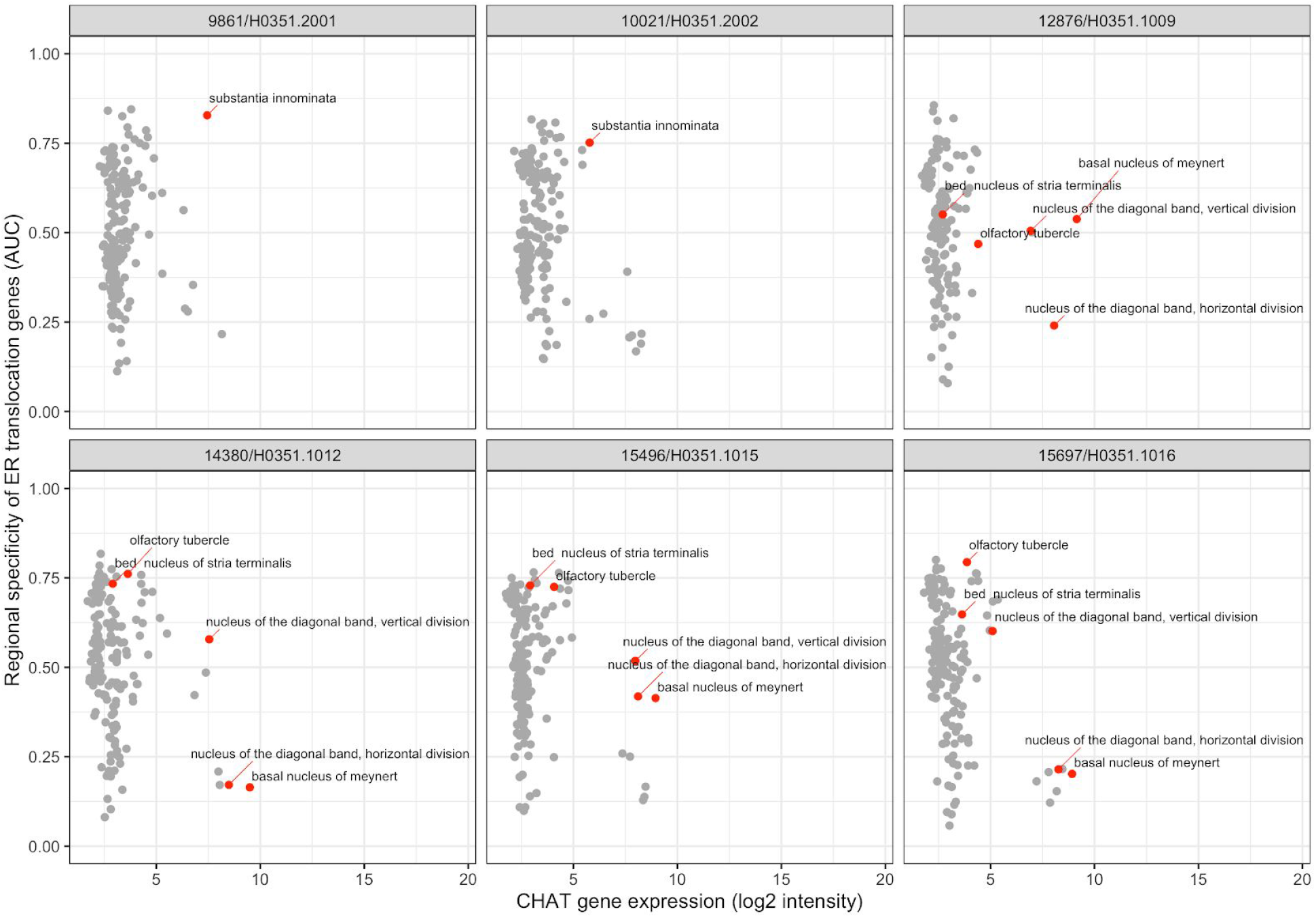
Scatter plots showing specific enrichment for ER translocation genes (y-axis) and Choline O-Acetyltransferase (*CHAT*) gene expression on the x-axis in each brain. Each point is a brain region with red marking the substantia innominata and its nuclei.

### ER translocation genes are highly expressed in mouse hypothalamic and cholinergic neurons

Mirroring our analyses of *P. gingivalis* associated genes, we next characterized the expression patterns of the ER translocation genes at a finer resolution in the mouse nervous system. We observe that the 79 mouse homologs of the ER translocation genes are not evenly expressed across clusters in this mouse single-cell atlas. The top 20 most enriched transcriptomic cell types clusters are provided in Table 4 (full listing in Supplement Table 9). Four cholinergic enteric neuron clusters are within the top ten cell clusters. Of the 14 cholinergic clusters in this atlas, 12 are enriched for ER translocation gene expression (all AUC > 0.5 and p_FDR_ < 0.05). Relative to the 265 tested cell type clusters, the cholinergic groups are enriched for higher AUC values (*p* < 0.0001, Mann-Whitney U test). The top-ranked cholinergic cell clusters from brain tissue are listed as telencephalon interneurons with an annotated location of striatum and amygdala (TECHO, AUC = 0.822, ranked 21). In Table 4, it is also clear that hypothalamic cells strongly express the ER translocation genes. Of the 14 total hypothalamic cell clusters, 7 are enriched for ER translocation gene expression (AUC > 0.57, p_FDR_ < 0.05). All of the hypothalamic cell type clusters enriched for ER translocation genes are peptidergic (or produce peptide hormone precursors). Three of the top hypothalamic transcriptomic cell types were annotated as having probable neuroanatomical locations that are in or near the basal forebrain. Specifically, in addition to hypothalamic regions, the HYPEP5 cluster lists the diagonal band nucleus and the HYPEP8 cluster names the medial septal nucleus as probable locations of those clustered cells (Zeisel et al. 2018). While substantia innominata is not mentioned in this atlas, both the mouse and human regional analyses highlight the basal forebrain, hypothalamus and cholinergic system.

**Table 4.**
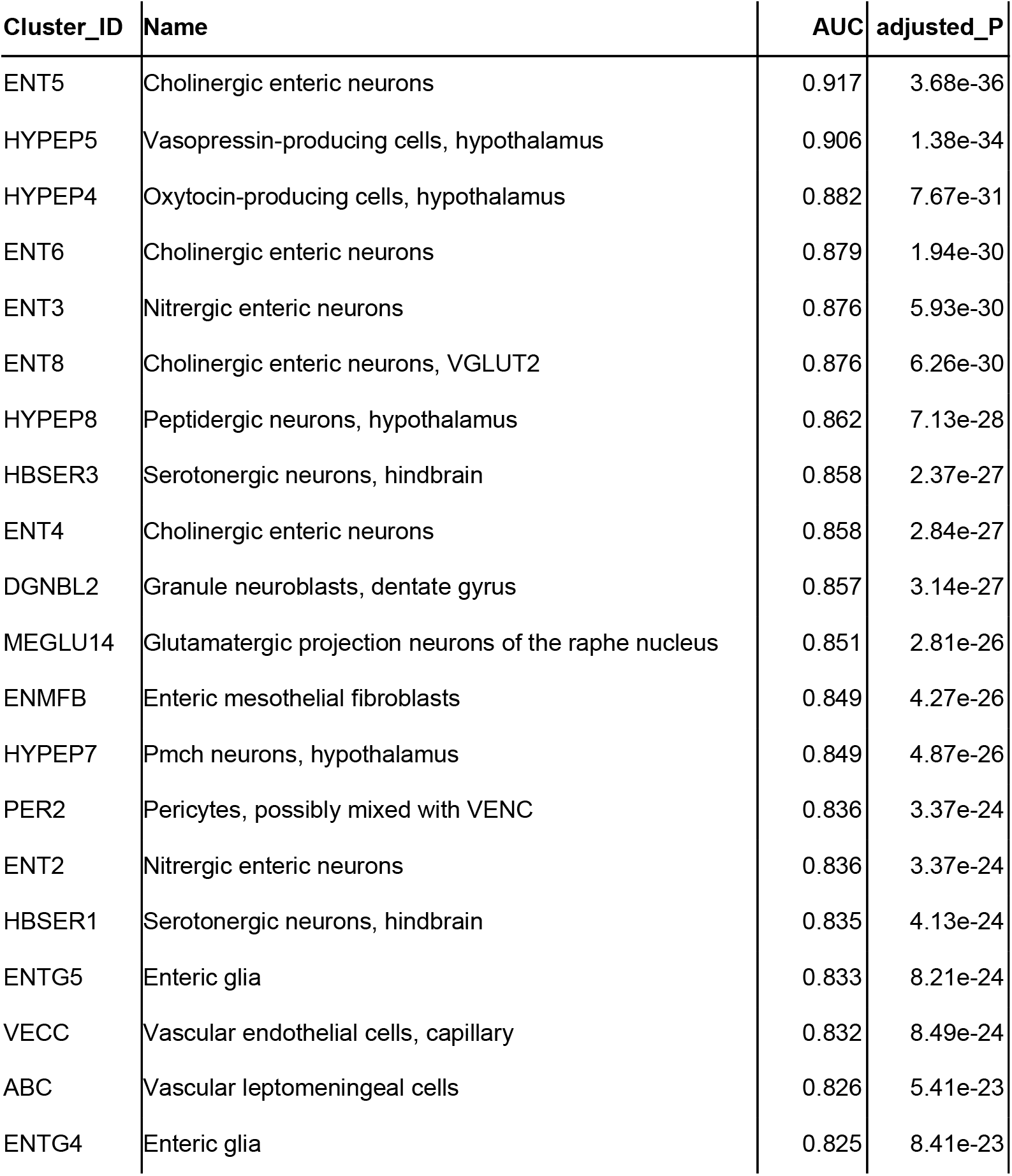
ER translocation gene enrichment statistics for cholinergic transcriptomic cell type clusters.

### High neuroanatomical convergence of *P. gingivalis* associated genes

We next tested for overlap between the two brain-wide patterns derived from separate sources of *P. gingivalis* associated genes. Within the top ten regions, the central gray of the pons, septal nuclei, and substantia innominata appear in both enrichment results. More specifically, the anterior hypothalamic area, the most enriched region for the genes associated with *P. gingivalis* reads, is the top-ranked hypothalamic region for the ER translocation results (ranked 51 of 232 overall, AUC = 0.697, p_FDR_ < 10^−10^). Conversely, the substantia innominata, which is most enriched for the ER translocation genes, is ranked 4th for genes associated with *P. gingivalis* reads. This spatial convergence within the top hits extends brain-wide with 48 brain regions overlapping between two lists of significantly enriched structures (hypergeometric test, p < 10^−6^) and correlation coefficient of AUC values of 0.36 (p < 10^−7^). This brain-wide spatial agreement is visualized in Figures 4 and 5. As shown in Figure 5B, high agreement is also observed in the single-cell atlas of the mouse nervous system (r = 0.72, p < 10^−43^). In particular, enriched expression in cholinergic neurons is evident for both sets of *P. gingivalis* associated genes. In summary, genes associated with gingipain susceptibility and *P. gingivalis* presence are highly expressed in hypothalamic, cholinergic neurons, and basal forebrain regions.

**Figure 4.**
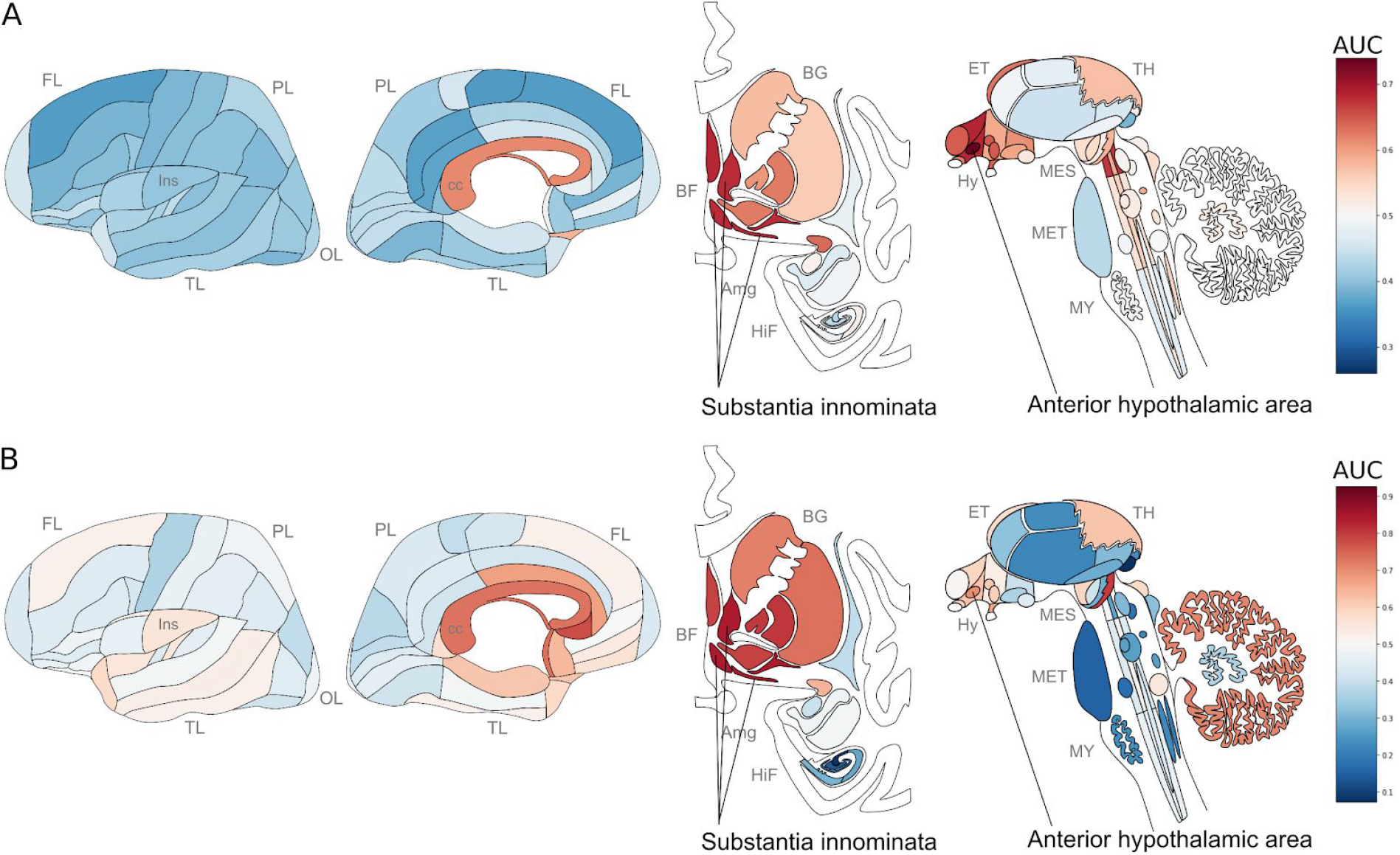
Neuroanatomical heatmaps marking specific expression of the genes up-regulated in samples with *P. gingivalis* reads (A) and ER translocation genes (B). AUC values range from depleted expression in dark blue to enriched in dark red. Brain region abbreviations: frontal lobe (FL); parietal lobe (PL); temporal lobe (TL); occipital lobe (OL); basal forebrain (BF); basal ganglia (BG); amygdala (AmG); hippocampal formation (HiF); epithalamus (EP); thalamus (TH); hypothalamus (Hy); mesencephalon (MES); metencephalon (MET) and myelencephalon (MY). Anatomical template images are from the Allen Human Brain Reference Atlas (S.-L. Ding et al. 2016).

**Figure 5.**
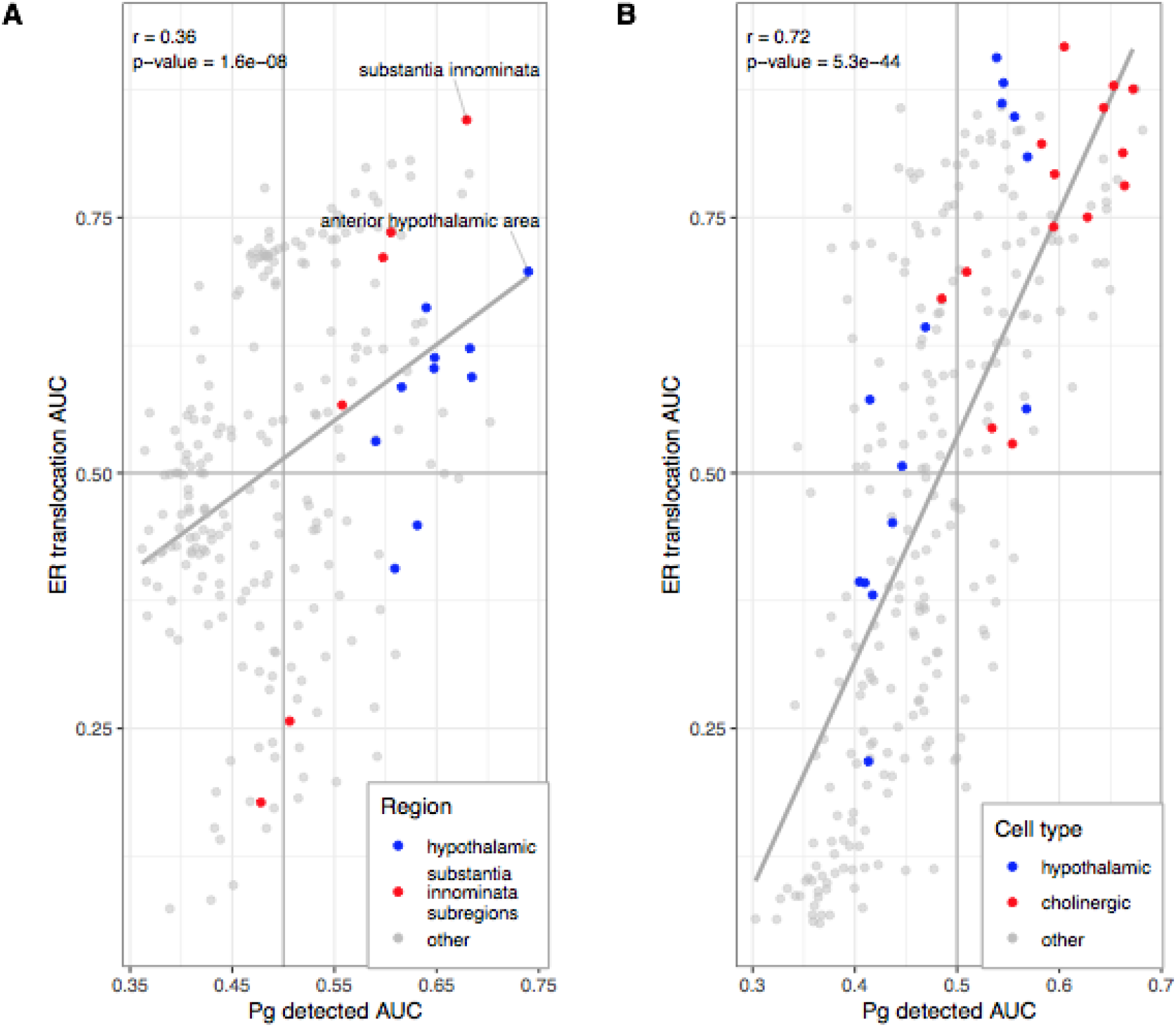
Scatterplots comparing anatomical enrichment of ER translocation genes (y-axis) and up-regulated in samples with *P. gingivalis* reads (x-axis). AUC values are plotted to show enrichment in the human brain atlas (A) and the mouse nervous system (B). Hypothalamic regions and transcriptomic cell type clusters are marked in blue. Cholinergic clusters, substantia innominata and it’s subregions are marked in red. Fitted linear regression lines for all points are shown in grey.

## Discussion

In this study, we characterized protein susceptibility to gingipain cleavage and genes differentially expressed in brain tissue with *P. gingivalis*. As expected, we found that genes with high arginine and lysine proportions are enriched in proteins that bind RNA and DNA. We focused on a specific set of these genes that participate in ER translocation and have previously been shown to be up-regulated in periodontitis and Alzheimer’s disease (Lundmark et al. 2018; Patel et al. 2020; Felsky et al. 2020). We directly link these findings to *P. gingivalis* by showing that these genes are also up-regulated in brain tissue with detected *P. gingivalis* RNA. This convergence between proteins susceptible to gingipain cleavage and the transcriptomic response to *P. gingivalis* motivated our neuroanatomical characterization of these genes. In this spatial analysis, we again observe agreement, with enrichment for cholinergic neurons, basal forebrain and hypothalamic regions. Regions near ventricles and peripheral neurons are also enriched, suggesting relevance to *P. gingivalis* brain entry. While gingipain levels have been shown to correlate with tau and amyloid pathology (Dominy et al. 2019; Ilievski et al. 2018; Poole et al. 2015), we are the first to associate this virulence factor with the cholinergic hypothesis of Alzheimer’s disease.

Several studies have examined arginine metabolism in the context of Alzheimer’s disease. Downstream metabolites of arginine are involved in microtubule assembly and stabilization (Wu and Morris 1998). For example, nitric oxide synthase (NOS) occurs in neuronal and endothelial cells, where it converts L-arginine to L-citrulline and nitric oxide gas. NOS is crucial for maintaining cerebral blood flow, synaptic plasticity, learning and memory (Esplugues 2002). Impairments in these areas are seen in Alzheimer’s disease. Disruption of arginine metabolism was found in Alzheimer’s disease brains (P. Liu et al. 2014) and in plasma and cerebrospinal fluid of individuals diagnosed with mild cognitive impairment and Alzheimer’s disease (Trushina et al. 2013). Accordingly, several experimental studies have detailed altered arginine metabolism in human cell lines and mouse models of Alzheimer’s disease (Vemula et al. 2019; Jko et al. 2016; Bergin et al. 2018; Kan et al. 2015). Lysine has also been studied in the context of Alzheimer’s disease with limited results (Griffin and Bradshaw 2017). These studies and our findings suggest disruptions of arginine metabolism should be examined for associations with gingipain activity in Alzheimer’s disease.

In our postmortem brain tissue analyses, several differentially expressed genes are related to amyloid processing and inflammation. For example, *ZNF267*, the 9th most up-regulated gene, was selected as one of the ten genes used in a blood-based transcriptomic panel for diagnosis Alzheimer’s Disease (Fehlbaum-Beurdeley et al. 2012). The most significantly up-regulated gene, integral membrane protein 2B (*ITM2B*, pFDR < 0.0011), regulates processing of amyloid-beta and inhibits amyloid aggregation (Kim et al. 2008). Mutations in this gene that encodes the BRI2 protein that causes familial British and Danish dementia (Vidal et al. 1999, 2000). We also note that the 12th most downregulated gene, Myc associated zinc finger protein (*MAZ*), is also known as serum amyloid A-activating factor-1 (SAF-1) due to its role in regulating serum amyloid A in response to inflammation (A. Ray and Ray 1996). Expression of the mouse homolog MAZ is increased in an Alzheimer’s disease mouse model (L. Liu et al. 2017), and murine overexpression increases the risk of severe arthritis (Alpana Ray et al. 2004). Like *P. gingivalis*, serum amyloid A is suspected to be involved in the pathogenesis of arthritis, atherosclerosis, amyloidosis and Alzheimer’s disease (Kindy et al. 1999; Alpana Ray et al. 2006; Getz, Krishack, and Reardon 2016; Targońska-Stępniak and Majdan 2014). Lastly, the 3rd most down-regulated gene, *DHX37*, harbours a rare frameshift mutation that segregates with Alzheimer’s disease in one family (Zhang et al. 2019). While our differential expression analysis highlighted ER translocation genes, several others link amyloid processing and diseases associated with *P. gingivalis*.

In regards to neuroinflammation, several chemokine ligands are up-regulated in brain samples with *P. gingivalis* reads. For example, CXCL1 and it’s paralog CXCL2 are ranked 25th and 336th genome-wide, respectively (both pFDR < 0.001). CXCL1 is known to function as a chemoattractant for neutrophils (Sawant et al. 2016). Also, CXCL2 is triggered by inflammation, and its expression is induced by *P. gingivalis* and in mouse models of periodontitis (Zhou et al. 2005; Tsujimoto et al. 2005; Kawagishi et al. 2001). Although our cell type-specific analyses did not highlight microglia, the ER translocation genes have been shown to progressively increase in expression across in AD-associated states of microglial activation (Patel et al. 2020). Overall, we did not observe strong neuroinflammatory signals. This may be due to our use of neuropathologically normal brains and the ability of *P. gingivalis* to selectively induce inflammatory responses (Darveau, Hajishengallis, and Curtis 2012).

Genes that function in cell-cell contact were down-regulated in brain tissue with detected *P. gingivalis*. Specifically, the top two GO groups enriched for down-regulation were “homophilic cell adhesion via plasma membrane adhesion molecules” and “regulation of synaptic plasticity”. These are independent signals as only two genes are in both of these sets. A recent study by Haditsch et al., found that neurons derived from human inducible pluripotent stem cells, when infected with *P. gingivalis*, had a significant loss of synapse density (Haditsch et al. 2020a). This loss was more pronounced than neuron cell death. In agreement, our results do not suggest differences in estimated neuron proportions. It is known that synapse loss is an early and significant event in Alzheimer’s disease (Scheff et al. 2015). In addition, in the aging and Alzheimer’s brain, decreases in synapse number and gene expression are observed but not neuron counts (French et al. 2017; Mostany et al. 2013). Haditsch and colleagues suggest synapse loss is due to microtubule destabilization caused by tau degradation by gingipains. In support, the most differentially expressed gene in the cell adhesion GO group is Microtubule Associated Protein Tau (MAPT, pFDR < 0.005, 198th most down-regulated gene). However, our results suggest that transcriptional responses may partially cause reduced synapse density and cell adhesion. Cell-to-cell contact has been shown to increase *P. gingivalis* transmission rate (Li et al. 2008), suggesting that these transcriptional responses may reduce *P. gingivalis* persistence.

The ribosome, RNA processing, and protein synthesis have been previously associated with mild cognitive impairment, atherosclerosis, and Alzheimer’s disease (Q. Ding et al. 2005; Sajdel-Sulkowska and Marotta 1984; Langstrom et al. 1989; Hernández-Ortega et al. 2016; Nyhus et al. 2019). In agreement with gingipain susceptibility, a proteomic study found a lower abundance of RNA splicing proteins in cerebral atherosclerosis (Wingo et al. 2020). Dysregulated splicing is also demonstrated by increased intron retention in Alzheimer’s disease (Adusumalli et al. 2019). Similarly, RNA quality, which is measured from primarily ribosomal RNA is lower in brain tissue from dementia cases (Miller et al. 2017). Pathological tau, a key marker of Alzheimer’s disease, has been shown to determine translational selectivity and co-localize with ribosomes (Koren et al. 2019; Meier et al. 2016). Also, major histone acetylation differences have been associated with tau pathology (Klein et al. 2019). These findings of disruptions in processes involving nucleic acid interactions, match the functions of proteins enriched for arginine and lysine residues.

While proteins that interact with RNA and DNA are broadly enriched for arginine and lysine, protein targeting to the ER is the top GO group enriched for up-regulation in samples with detected *P. gingivalis*. While speculative, the ER translocation genes point to mechanisms that support the gingipain hypothesis of Alzheimer’s disease. The gingipain hypothesis proposes that *P. gingivalis* infection of brain tissue causes Alzheimer’s disease (Dominy et al. 2019). In infected human cells, *P. gingivalis* is found in vacuoles that contain undegraded ribosomes (Dorn, Dunn, and Progulske-Fox 2001). *P. gingivalis* ferments amino acids as an energy source (Nelson et al. 2003). Co-localized ribosomes may provide a particularly digestible source of amino acids because of their enrichment for the positively charged residues that gingipains cleave. Cytosolically free *P. gingivalis* quickly localizes to the rough ER to form autophagosome-like vacuoles (Lee et al. 2018). However, it is not known how *P. gingivalis* traffics to the ER (Bélanger et al. 2006) and we speculate that ER translocation processes may be exploited by *P. gingivalis* to reach the endoplasmic reticulum. Sequestration of ribosomes in autophagosome-like vacuoles by *P. gingivalis* may cause increased transcription of cytosolic ribosomal protein genes because ribosome biogenesis is highly regulated (Lempiäinen and Shore 2009). Further study is needed to experimentally test if *P. gingivalis* exploits cytosolic ribosomes and ER translocation pathways.

The anterior hypothalamic area, oxytocin- and vasopressin-expressing neurons were strongly enriched for our *P. gingivalis* associated genes. The hypothalamus releases neuropeptides and peptide hormones that go through extensive pre-processing in the ER. Evidence of hypothalamic dysfunction and decreases in hypothalamic volume have been found in Alzheimer’s disease patients [reviewed in (Ishii and Iadecola 2015; Hiller and Ishii 2018)]. Specifically, significant deficits in sleep and circadian rhythm have been reported with associations to vasopressin in human studies (Hu et al. 2013; Harper et al. 2008). Orexin, which plays a major role in the sleep-wake cycle, has been associated with amyloid pathology in mouse models (Kang et al. 2009; Roh et al. 2014).

In Alzheimer’s patients, orexin levels in cerebrospinal fluid were correlated with amyloid-β42, sleep disruption and fragmentation (Gabelle et al. 2017; Liguori et al. 2016). In addition, galanin, which is strongly expressed in the hypothalamic neurons enriched in our analysis, is associated with sleep fragmentation in Alzheimer’s disease (Lim et al. 2014). Hypothalamic dysfunction in Alzheimer’s disease and its enrichment for *P. gingivalis* associated genes suggest this region is relevant to the gingipain hypothesis.

Our findings associate the substantia innominata and cholinergic neurons with *P. gingivalis* transcriptomic response and gingipain susceptibility. Enrichment in the medial habenula, a region with high expression of nicotinic cholinergic receptors, is also observed (Le Foll and French 2018). Extensive loss of cholinergic neurons in the substantia innominata has been found in Alzheimer’s patients (Whitehouse et al. 1982, 1981). This finding and many others form the basis of the cholinergic hypothesis of AD, which proposes that degeneration of cholinergic neurons in the basal forebrain substantially contributes to cognitive decline in AD patients (Hampel et al. 2019). This subcortical degeneration is thought to occur early in the disease process (M. Mesulam et al. 2004). In the context of the gingipain hypothesis, we note that reduced basal forebrain volume and cholinergic function follow after the removal of teeth in rodents (Kato et al. 1997; Onozuka et al. 2002; Avivi-Arber et al. 2016) and oral acetylcholine levels are correlated with periodontal disease severity (Apatzidou et al. 2018). High expression of the ER translocation genes in cholinergic neurons may be required to support acetylcholinesterase processing in the ER (Dobbertin et al. 2009). In the mouse atlas, cholinergic neurons in the enteric nervous system had the highest expression of the ER translocation genes. In the context of AD, loss of enteric cholinergic neurons has been observed in a transgenic mouse model of the disease (Han et al. 2017). Taken together, our *P. gingivalis* associated genes highlight cholinergic neurons and the substantia innominata, providing an anatomical between the gingipain and cholinergic hypotheses.

## Conclusion

Proteins that are enriched for arginine and lysine residues, which are potential gingipain cleavage sites bind RNA and DNA. While ribosomal genes are enriched for these residues, a specific set which function in SRP dependent ER cotranslational translocation has been previously associated with periodontitis and Alzheimer’s disease (Patel et al. 2020; Felsky et al. 2020). These genes are also up-regulated in brain tissue with detected *P. gingivalis* RNA, suggesting a transcriptional response to gingipain driven degradation of ER localized ribosomes. Neuroanatomical analysis of the *P. gingivalis* associated genes marks cholinergic neurons, basal forebrain and hypothalamic regions. These results link the gingipain and cholinergic hypotheses. At the mechanistic level, the highlighted translocation genes may explain how *P. gingivalis* transits to the ER upon cell entry. Also, down-regulation of synapse and cell-cell contact genes support findings of synapse loss upon *P. gingivalis* infection (Haditsch et al. 2020b). In summary, our findings detail *P. gingivalis* response and susceptibility at the molecular and anatomical levels that suggest new associations that are relevant to Alzheimer’s disease pathogenesis.

## Acknowledgements

We thank Navona Calarco for help with the analysis of the single-cell atlas of the mouse nervous system. We thank the Allen Institute for Brain Science for creating the transcriptomic atlas of the human brain. We thank Ed Lein, Michael Hawrylycz, Jeremy Miller, Taylor Schmitz, Shreejoy Tripathy, Karina Carneiro, and Daniel Felsky for their insightful comments and suggestions.

This study was supported by the CAMH Foundation, CAMH Discovery Fund, and a National Science and Engineering Research Council of Canada (NSERC) Discovery Grant to LF.

## References

Adusumalli, Swarnaseetha, Zhen-Kai Ngian, Wei-Qi Lin, Touati Benoukraf, and Chin-Tong Ong. 2019. “Increased Intron Retention Is a Post-Transcriptional Signature Associated with Progressive Aging and Alzheimer’s Disease.” Aging Cell 18 (3): e12928.

Ahn, Jiyoung, Stephanie Segers, and Richard B. Hayes. 2012. “Periodontal Disease, Porphyromonas Gingivalis Serum Antibody Levels and Orodigestive Cancer Mortality.” Carcinogenesis 33 (5): 1055–58.

Apatzidou, Danae A., Achilleas Iskas, Antonis Konstantinidis, Abeer M. Alghamdi, Maria Tumelty, David F. Lappin, and Christopher J. Nile. 2018. “Clinical Associations between Acetylcholine Levels and Cholinesterase Activity in Saliva and Gingival Crevicular Fluid and Periodontal Diseases.” Journal of Clinical Periodontology 45 (10): 1173–83.

Ashburner, M., C. A. Ball, J. A. Blake, D. Botstein, H. Butler, J. M. Cherry, A. P. Davis, et al. 2000. “Gene Ontology: Tool for the Unification of Biology. The Gene Ontology Consortium.” Nature Genetics 25 (1): 25–29.

Avivi-Arber, Limor, Ze ‘ev Seltzer, Miriam Friedel, Jason P. Lerch, Massieh Moayedi, Karen D. Davis, and Barry J. Sessle. 2016. “Widespread Volumetric Brain Changes Following Tooth Loss in Female Mice.” Frontiers in Neuroanatomy 10: 121.

Baker, N. A., D. Sept, S. Joseph, M. J. Holst, and J. A. McCammon. 2001. “Electrostatics of Nanosystems: Application to Microtubules and the Ribosome.” Proceedings of the National Academy of Sciences of the United States of America 98 (18): 10037–41.

Bélanger, Myriam, Paulo H. Rodrigues, William A. Dunn Jr, and Ann Progulske-Fox. 2006. “Autophagy: A Highway for Porphyromonas Gingivalis in Endothelial Cells.” Autophagy 2 (3): 165–70.

Bergin, D. H., Y. Jing, B. G. Mockett, H. Zhang, W. C. Abraham, and P. Liu. 2018. “Altered Plasma Arginine Metabolome Precedes Behavioural and Brain Arginine Metabolomic Profile Changes in the APPswe/PS1ΔE9 Mouse Model of Alzheimer’s Disease.” Translational Psychiatry 8 (1): 108.

Bingham, Clifton O., 3rd, and Malini Moni. 2013. “Periodontal Disease and Rheumatoid Arthritis: The Evidence Accumulates for Complex Pathobiologic Interactions.” Current Opinion in Rheumatology 25 (3): 345–53.

Birse, D. E., U. Kapp, K. Strub, S. Cusack, and A. Aberg. 1997. “The Crystal Structure of the Signal Recognition Particle Alu RNA Binding Heterodimer, SRP9/14.” The EMBO Journal 16 (13): 3757–66.

Blanco, Celia, Marco Bayas, Fu Yan, and Irene A. Chen. 2018. “Analysis of Evolutionarily Independent Protein-RNA Complexes Yields a Criterion to Evaluate the Relevance of Prebiotic Scenarios.” Current Biology: CB 28 (4): 526–37.e5.

Bostanci, Nagihan, and Georgios N. Belibasakis. 2012. “Porphyromonas Gingivalis: An Invasive and Evasive Opportunistic Oral Pathogen.” FEMS Microbiology Letters 333 (1): 1–9.

Braak, H., and E. Braak. 1991. “Neuropathological Stageing of Alzheimer-Related Changes.” Acta Neuropathologica 82 (4): 239–59.

Bui, Fiona Q., Cassio Luiz Coutinho Almeida-da-Silva, Brandon Huynh, Alston Trinh, Jessica Liu, Jacob Woodward, Homer Asadi, and David M. Ojcius. 2019. “Association between Periodontal Pathogens and Systemic Disease.” Biomedical Journal 42 (1): 27–35.

Carlson, Marc. 2016a. “GO.db: A Set of Annotation Maps Describing the Entire Gene Ontology.”

Carlson, Marc. 2016b. “org.Hs.eg.db: Genome Wide Annotation for Human.”

Darmanis, Spyros, Steven A. Sloan, Ye Zhang, Martin Enge, Christine Caneda, Lawrence M. Shuer, Melanie G. Hayden Gephart, Ben A. Barres, and Stephen R. Quake. 2015. “A Survey of Human Brain Transcriptome Diversity at the Single Cell Level.” Proceedings of the National Academy of Sciences of the United States of America 112 (23): 7285–90.

Darveau, R. P., G. Hajishengallis, and M. A. Curtis. 2012. “Porphyromonas Gingivalis as a Potential Community Activist for Disease.” Journal of Dental Research 91 (9): 816–20.

DeKosky, S. T., R. E. Harbaugh, F. A. Schmitt, R. A. Bakay, H. C. Chui, D. S. Knopman, T. M. Reeder, A. G. Shetter, H. J. Senter, and W. R. Markesbery. 1992. “Cortical Biopsy in Alzheimer’s Disease: Diagnostic Accuracy and Neurochemical, Neuropathological, and Cognitive Correlations. Intraventricular Bethanecol Study Group.” Annals of Neurology 32 (5): 625–32.

Diego, Iñaki de, Iñaki de Diego, Florian Veillard, Maryta N. Sztukowska, Tibisay Guevara, Barbara Potempa, Anja Pomowski, James A. Huntington, Jan Potempa, and F. Xavier Gomis-Rüth. 2014. “Structure and Mechanism of Cysteine Peptidase Gingipain K (Kgp), a Major Virulence Factor ofPorphyromonas Gingivalisin Periodontitis.” Journal of Biological Chemistry. https://doi.org/10.1074/jbc.m114.602052.

Ding, Qunxing, William R. Markesbery, Qinghua Chen, Feng Li, and Jeffrey N. Keller. 2005. “Ribosome Dysfunction Is an Early Event in Alzheimer’s Disease.” The Journal of Neuroscience: The Official Journal of the Society for Neuroscience 25 (40): 9171–75.

Ding, Song-Lin, Joshua J. Royall, Susan M. Sunkin, Lydia Ng, Benjamin A. C. Facer, Phil Lesnar, Angie Guillozet-Bongaarts, et al. 2016. “Comprehensive Cellular-Resolution Atlas of the Adult Human Brain.” The Journal of Comparative Neurology 524 (16): 3127–3481.

Dobbertin, Alexandre, Anna Hrabovska, Korami Dembele, Shelley Camp, Palmer Taylor, Eric Krejci, and Véronique Bernard. 2009. “Targeting of Acetylcholinesterase in Neurons in Vivo: A Dual Processing Function for the Proline-Rich Membrane Anchor Subunit and the Attachment Domain on the Catalytic Subunit.” The Journal of Neuroscience: The Official Journal of the Society for Neuroscience 29 (14): 4519–30.

Dominy, Stephen S., Casey Lynch, Florian Ermini, Malgorzata Benedyk, Agata Marczyk, Andrei Konradi, Mai Nguyen, et al. 2019. “Porphyromonas Gingivalis in Alzheimer’s Disease Brains: Evidence for Disease Causation and Treatment with Small-Molecule Inhibitors.” Science Advances 5 (1): eaau3333.

Dorn, B. R., W. A. Dunn Jr, and A. Progulske-Fox. 2001. “Porphyromonas Gingivalis Traffics to Autophagosomes in Human Coronary Artery Endothelial Cells.” Infection and Immunity 69 (9): 5698–5708.

Dumitriu, Alexandra, Javad Golji, Adam T. Labadorf, Benbo Gao, Thomas G. Beach, Richard H. Myers, Kenneth A. Longo, and Jeanne C. Latourelle. 2016. “Integrative Analyses of Proteomics and RNA Transcriptomics Implicate Mitochondrial Processes, Protein Folding Pathways and GWAS Loci in Parkinson Disease.” BMC Medical Genomics 9 (January): 5.

Esplugues, Juan V. 2002. “NO as a Signalling Molecule in the Nervous System.” British Journal of Pharmacology 135 (5): 1079.

Farquharson, D., J. P. Butcher, and S. Culshaw. 2012. “Periodontitis, Porphyromonas, and the Pathogenesis of Rheumatoid Arthritis.” Mucosal Immunology 5 (2): 112–20.

Felsky, Daniel, Sanjeev Sariya, Ismael Santa-Maria, Julie A. Schneider, David A. Bennett, Richard Mayeux, Philip L. De Jager, and Giuseppe Tosto. 2020. “The Caribbean-Hispanic Alzheimer’s Brain Transcriptome Reveals Ancestry-Specific Disease Mechanisms.” bioRxiv. https://doi.org/10.1101/2020.05.28.122234.

Fiorillo, Luca, Gabriele Cervino, Luigi Laino, Cesare D’Amico, Rodolfo Mauceri, Tolga Fikret Tozum, Michele Gaeta, and Marco Cicciù. 2019. “Porphyromonas Gingivalis, Periodontal and Systemic Implications: A Systematic Review.” Dental Journal 7 (4). https://doi.org/10.3390/dj7040114.

Frankish, Adam, Mark Diekhans, Anne-Maud Ferreira, Rory Johnson, Irwin Jungreis, Jane Loveland, Jonathan M. Mudge, et al. 2019. “GENCODE Reference Annotation for the Human and Mouse Genomes.” Nucleic Acids Research 47 (D1): D766–73.

French, Leon, Tianzhou Ma, Hyunjung Oh, George C. Tseng, and Etienne Sibille. 2017. “Age-Related Gene Expression in the Frontal Cortex Suggests Synaptic Function Changes in Specific Inhibitory Neuron Subtypes.” Frontiers in Aging Neuroscience 9 (May): 162.

Gabelle, Audrey, Isabelle Jaussent, Christophe Hirtz, Jérôme Vialaret, Sophie Navucet, Caroline Grasselli, Philippe Robert, Sylvain Lehmann, and Yves Dauvilliers. 2017. “Cerebrospinal Fluid Levels of Orexin-A and Histamine, and Sleep Profile within the Alzheimer Process.” Neurobiology of Aging 53 (May): 59–66.

Getz, Godfrey S., Paulette A. Krishack, and Catherine A. Reardon. 2016. “Serum Amyloid A and Atherosclerosis.” Current Opinion in Lipidology 27 (5): 531–35.

Griffin, Jeddidiah W. D., and Patrick C. Bradshaw. 2017. “Amino Acid Catabolism in Alzheimer’s Disease Brain: Friend or Foe?” Oxidative Medicine and Cellular Longevity 2017 (February): 5472792.

Guo, Yonghua, Ky-Anh Nguyen, and Jan Potempa. 2010. “Dichotomy of Gingipains Action as Virulence Factors: From Cleaving Substrates with the Precision of a Surgeon’s Knife to a Meat Chopper-like Brutal Degradation of Proteins.” Periodontology 2000 54 (1): 15–44.

Haditsch, Ursula, Theresa Roth, Leo Rodriguez, Sandy Hancock, Thomas Cecere, Mai Nguyen, Shirin Arastu-Kapur, et al. 2020a. “Alzheimer’s Disease-Like Neurodegeneration in Porphyromonas Gingivalis Infected Neurons with Persistent Expression of Active Gingipains.” Journal of Alzheimer’s Disease. https://doi.org/10.3233/jad-200393.

Shirin Arastu-Kapur, et al. 2020b. “Alzheimer’s Disease-Like Neurodegeneration in Porphyromonas Gingivalis Infected Neurons with Persistent Expression of Active Gingipains.” Journal of Alzheimer’s Disease: JAD 75 (4): 1361–76.

Hampel, Harald, M-M Mesulam, A. C. Cuello, A. S. Khachaturian, A. Vergallo, M. R. Farlow, P. J. Snyder, E. Giacobini, Z. S. Khachaturian, and Cholinergic System Working Group, and for the Alzheimer Precision Medicine Initiative (APMI). 2019. “Revisiting the Cholinergic Hypothesis in Alzheimer’s Disease: Emerging Evidence from Translational and Clinical Research.” The Journal of Prevention of Alzheimer’s Disease 6 (1): 2–15.

Han, Xiaolei, Shi Tang, Lingling Dong, Lin Song, Yi Dong, Yongxiang Wang, and Yifeng Du. 2017. “Loss of Nitrergic and Cholinergic Neurons in the Enteric Nervous System of APP/PS1 Transgenic Mouse Model.” Neuroscience Letters 642 (March): 59–65.

Haraszthy, V. I., J. J. Zambon, M. Trevisan, M. Zeid, and R. J. Genco. 2000. “Identification of Periodontal Pathogens in Atheromatous Plaques.” Journal of Periodontology. https://doi.org/10.1902/jop.2000.71.10.1554.

Harper, David G., Edward G. Stopa, Victoria Kuo-Leblanc, Ann C. McKee, Kentaro Asayama, Ladislav Volicer, Neil Kowall, and Andrew Satlin. 2008. “Dorsomedial SCN Neuronal Subpopulations Subserve Different Functions in Human Dementia.” Brain: A Journal of Neurology 131 (Pt 6): 1609–17.

Hawrylycz, Michael J., Ed S. Lein, Angela L. Guillozet-Bongaarts, Elaine H. Shen, Lydia Ng, Jeremy A. Miller, Louie N. van de Lagemaat, et al. 2012. “An Anatomically Comprehensive Atlas of the Adult Human Brain Transcriptome.” Nature 489 (7416): 391–99.

Hernandez-Ortega, Karina, Paula Garcia-Esparcia, Laura Gil, José J. Lucas, and Isidre Ferrer. 2016. “Altered Machinery of Protein Synthesis in Alzheimer’s: From the Nucleolus to the Ribosome.” Brain Pathology 26 (5): 593–605.

Hiller, Abigail J., and Makoto Ishii. 2018. “Disorders of Body Weight, Sleep and Circadian Rhythm as Manifestations of Hypothalamic Dysfunction in Alzheimer’s Disease.” Frontiers in Cellular Neuroscience 12 (December): 471.

Hu, Kun, David G. Harper, Steven A. Shea, Edward G. Stopa, and Frank A. J. L. Scheer. 2013. “Noninvasive Fractal Biomarker of Clock Neurotransmitter Disturbance in Humans with Dementia.” Scientific Reports 3: 2229.

Ilievski, Vladimir, Paulina K. Zuchowska, Stefan J. Green, Peter T. Toth, Michael E. Ragozzino, Khuong Le, Haider W. Aljewari, Neil M. O’Brien-Simpson, Eric C. Reynolds, and Keiko Watanabe. 2018. “Chronic Oral Application of a Periodontal Pathogen Results in Brain Inflammation, Neurodegeneration and Amyloid Beta Production in Wild Type Mice.” PloS One 13 (10): e0204941.

Ishida, Naoyuki, Yuichi Ishihara, Kazuto Ishida, Hiroyuki Tada, Yoshiko Funaki-Kato, Makoto Hagiwara, Taslima Ferdous, et al. 2017. “Periodontitis Induced by Bacterial Infection Exacerbates Features of Alzheimer’s Disease in Transgenic Mice.” NPJ Aging and Mechanisms of Disease 3 (November): 15.

Ishii, Makoto, and Costantino Iadecola. 2015. “Metabolic and Non-Cognitive Manifestations of Alzheimer’s Disease: The Hypothalamus as Both Culprit and Target of Pathology.” Cell Metabolism 22 (5): 761–76.

Jko, Henryk, Anna Wilkaniec, Magdalena Cielik, Wojciech Hilgier, Magdalena Gssowska, Walter J. Lukiw, and Agata Adamczyk. 2016. “Altered Arginine Metabolism in Cells Transfected with Human Wild-Type Beta Amyloid Precursor Protein (APP).” Current Alzheimer Research 13 (9): 1030–39.

Kang, Jae-Eun, Miranda M. Lim, Randall J. Bateman, James J. Lee, Liam P. Smyth, John R. Cirrito, Nobuhiro Fujiki, Seiji Nishino, and David M. Holtzman. 2009. “Amyloid-Beta Dynamics Are Regulated by Orexin and the Sleep-Wake Cycle.” Science 326 (5955): 1005–7.

Kan, Matthew J., Jennifer E. Lee, Joan G. Wilson, Angela L. Everhart, Candice M. Brown, Andrew N. Hoofnagle, Marilyn Jansen, Michael P. Vitek, Michael D. Gunn, and Carol A. Colton. 2015. “Arginine Deprivation and Immune Suppression in a Mouse Model of Alzheimer’s Disease.” The Journal of Neuroscience: The Official Journal of the Society for Neuroscience 35 (15): 5969–82.

Kato, T., T. Usami, Y. Noda, M. Hasegawa, M. Ueda, and T. Nabeshima. 1997. “The Effect of the Loss of Molar Teeth on Spatial Memory and Acetylcholine Release from the Parietal Cortex in Aged Rats.” Behavioural Brain Research 83 (1-2): 239–42.

Kawagishi, C., K. Kurosaka, N. Watanabe, and Y. Kobayashi. 2001. “Cytokine Production by Macrophages in Association with Phagocytosis of Etoposide-Treated P388 Cells in Vitro and in Vivo.” Biochimica et Biophysica Acta 1541 (3): 221–30.

Kellesarian, Sergio Varela, Michael Yunker, Hans Malmstrom, Khalid Almas, Georgios E. Romanos, and Fawad Javed. 2018. “Male Infertility and Dental Health Status: A Systematic Review.” American Journal of Men’s Health 12 (6): 1976–84.

Kim, Jungsu, Victor M. Miller, Yona Levites, Karen Jansen West, Craig W. Zwizinski, Brenda D. Moore, Fredrick J. Troendle, et al. 2008. “BRI2 (ITM2b) Inhibits Abeta Deposition in Vivo.” The Journal of Neuroscience: The Official Journal of the Society for Neuroscience 28 (23): 6030–36.

Kindy, Mark S., Jin Yu, Jun-Tao Guo, and Hong Zhu. 1999. “Apolipoprotein Serum Amyloid A in Alzheimer’s Disease.” Journal of Alzheimer’s Disease: JAD 1 (3): 155–67.

Klareskog, Lars, Johan Rönnelid, Karin Lundberg, Leonid Padyukov, and Lars Alfredsson. 2008. “Immunity to Citrullinated Proteins in Rheumatoid Arthritis.” Annual Review of Immunology 26: 651–75.

Klein, Hans-Ulrich, Cristin McCabe, Elizabeta Gjoneska, Sarah E. Sullivan, Belinda J. Kaskow, Anna Tang, Robert V. Smith, et al. 2019. “Epigenome-Wide Study Uncovers Large-Scale Changes in Histone Acetylation Driven by Tau Pathology in Aging and Alzheimer’s Human Brains.” Nature Neuroscience 22 (1): 37–46.

Koren, Shon A., Matthew J. Hamm, Shelby E. Meier, Blaine E. Weiss, Grant K. Nation, Emad A. Chishti, Juan Pablo Arango, et al. 2019. “Tau Drives Translational Selectivity by Interacting with Ribosomal Proteins.” Acta Neuropathologica 137 (4): 571–83.

Langstrom, N. S., J. P. Anderson, H. G. Lindroos, B. Winblad, and W. C. Wallace. 1989. “Alzheimer’s Disease-Associated Reduction of Polysomal mRNA Translation.” Brain Research. Molecular Brain Research 5 (4): 259–69.

Lee, Kyulim, Joann S. Roberts, Chul Hee Choi, Kalina R. Atanasova, and Özlem Yilmaz. 2018. “Porphyromonas Gingivalis Traffics into Endoplasmic Reticulum-Rich-Autophagosomes for Successful Survival in Human Gingival Epithelial Cells.” Virulence 9 (1): 845–59.

Le Foll, Bernard, and Leon French. 2018. “Transcriptomic Characterization of the Human Habenula Highlights Drug Metabolism and the Neuroimmune System.” Frontiers in Neuroscience 12 (October): 742.

Leinonen, Rasko, Hideaki Sugawara, Martin Shumway, and International Nucleotide Sequence Database Collaboration. 2011. “The Sequence Read Archive.” Nucleic Acids Research 39 (Database issue): D19–21.

Lempiäinen, Harri, and David Shore. 2009. “Growth Control and Ribosome Biogenesis.” Current Opinion in Cell Biology 21 (6): 855–63.

Liguori, Claudio, Marzia Nuccetelli, Francesca Izzi, Giuseppe Sancesario, Andrea Romigi, Alessandro Martorana, Chiara Amoroso, et al. 2016. “Rapid Eye Movement Sleep Disruption and Sleep Fragmentation Are Associated with Increased Orexin-A Cerebrospinal-Fluid Levels in Mild Cognitive Impairment due to Alzheimer’s Disease.” Neurobiology of Aging 40 (April): 120–26.

Li, Ling, Raynald Michel, Joshua Cohen, Arthur Decarlo, and Emil Kozarov. 2008. “Intracellular Survival and Vascular Cell-to-Cell Transmission of Porphyromonas Gingivalis.” BMC Microbiology 8 (February): 26.

Lim, Andrew S. P., Brian A. Ellison, Joshua L. Wang, Lei Yu, Julie A. Schneider, Aron S. Buchman, David A. Bennett, and Clifford B. Saper. 2014. “Sleep Is Related to Neuron Numbers in the Ventrolateral Preoptic/intermediate Nucleus in Older Adults with and without Alzheimer’s Disease.” Brain: A Journal of Neurology 137 (Pt 10): 2847–61.

Liu, Chia-Chen, Chia-Chan Liu, Takahisa Kanekiyo, Huaxi Xu, and Guojun Bu. 2013. “Apolipoprotein E and Alzheimer Disease: Risk, Mechanisms and Therapy.” Nature Reviews. Neurology 9 (2): 106–18.

Liu, Lu, Yu-Jie Lai, Li-Ge Zhao, and Guo-Jun Chen. 2017. “Increased Expression of Myc-Interacting Zinc Finger Protein 1 in APP/PS1 Mice.” Experimental and Therapeutic Medicine 14 (6): 5751–56.

Liu, Ping, Michael S. Fleete, Yu Jing, Nicola D. Collie, Maurice A. Curtis, Henry J. Waldvogel, Richard L. M. Faull, Wickliffe C. Abraham, and Hu Zhang. 2014. “Altered Arginine Metabolism in Alzheimer’s Disease Brains.” Neurobiology of Aging 35 (9): 1992–2003.

Lönn, J., S. Ljunggren, K. Klarström-Engström, I. Demirel, T. Bengtsson, and H. Karlsson. 2018. “Lipoprotein Modifications by Gingipains ofPorphyromonas Gingivalis.” Journal of Periodontal Research. https://doi.org/10.1111/jre.12527.

Lott, Brittany Burton, Yongmei Wang, and Takuya Nakazato. 2013. “A Comparative Study of Ribosomal Proteins: Linkage between Amino Acid Distribution and Ribosomal Assembly.” BMC Biophysics 6 (1): 13.

Ludden, Christopher William. 2015. “Transcriptomic Evaluation of Peri-Implant Soft Tissue in Health and Disease.” The Ohio State University. https://etd.ohiolink.edu/!etd.send_file?accession=osu1435233979&disposition=inline.

Lundmark, Anna, Natalija Gerasimcik, Tove Båge, Anders Jemt, Annelie Mollbrink, Fredrik Salmén, Joakim Lundeberg, and Tülay Yucel-Lindberg. 2018. “Gene Expression Profiling of Periodontitis-Affected Gingival Tissue by Spatial Transcriptomics.” Scientific Reports 8 (1): 9370.

Meier, Shelby, Michelle Bell, Danielle N. Lyons, Jennifer Rodriguez-Rivera, Alexandria Ingram, Sarah N. Fontaine, Elizabeth Mechas, et al. 2016. “Pathological Tau Promotes Neuronal Damage by Impairing Ribosomal Function and Decreasing Protein Synthesis.” The Journal of Neuroscience: The Official Journal of the Society for Neuroscience 36 (3): 1001–7.

Mesulam, Marsel, Pamela Shaw, Deborah Mash, and Sandra Weintraub. 2004. “Cholinergic Nucleus Basalis Tauopathy Emerges Early in the Aging-MCI-AD Continuum.” Annals of Neurology 55 (6): 815–28.

Mesulam, M. M., E. J. Mufson, B. H. Wainer, and A. I. Levey. 1983. “Central Cholinergic Pathways in the Rat: An Overview Based on an Alternative Nomenclature (Ch1-Ch6).” Neuroscience 10 (4): 1185–1201.

Mikuls, Ted R., Jeffrey B. Payne, Richard A. Reinhardt, Geoffrey M. Thiele, Eileen Maziarz, Amy C. Cannella, V. Michael Holers, Kristine A. Kuhn, and James R. O’Dell. 2009. “Antibody Responses to Porphyromonas Gingivalis (P. Gingivalis) in Subjects with Rheumatoid Arthritis and Periodontitis.” International Immunopharmacology 9 (1): 38–42.

Miller, Jeremy A., Angela Guillozet-Bongaarts, Laura E. Gibbons, Nadia Postupna, Anne Renz, Allison E. Beller, Susan M. Sunkin, et al. 2017. “Neuropathological and Transcriptomic Characteristics of the Aged Brain.” eLife 6 (November). https://doi.org/10.7554/eLife.31126.

Miller, Jeremy A., Vilas Menon, Jeff Goldy, Ajamete Kaykas, Chang-Kyu Lee, Kimberly A. Smith, Elaine H. Shen, John W. Phillips, Ed S. Lein, and Mike J. Hawrylycz. 2014. “Improving Reliability and Absolute Quantification of Human Brain Microarray Data by Filtering and Scaling Probes Using RNA-Seq.” BMC Genomics 15 (February): 154.

Mostany, Ricardo, James E. Anstey, Kerensa L. Crump, Bohumil Maco, Graham Knott, and Carlos Portera-Cailliau. 2013. “Altered Synaptic Dynamics during Normal Brain Aging.” The Journal of Neuroscience: The Official Journal of the Society for Neuroscience 33 (9): 4094–4104.

Nadim, Rizwan, Jie Tang, Amena Dilmohamed, Siyang Yuan, Changhao Wu, Aishat T. Bakre, Martin Partridge, et al. 2020. “Influence of Periodontal Disease on Risk of Dementia: A Systematic Literature Review and a Meta-Analysis.” European Journal of Epidemiology, June. https://doi.org/10.1007/s10654-020-00648-x.

Nakano, Kazuhiko, Hiroaki Inaba, Ryota Nomura, Hirotoshi Nemoto, Munehiro Takeda, Hideo Yoshioka, Hajime Matsue, et al. 2006. “Detection of Cariogenic Streptococcus Mutans in Extirpated Heart Valve and Atheromatous Plaque Specimens.” Journal of Clinical Microbiology 44 (9): 3313–17.

Nelson, Karen E., Robert D. Fleischmann, Robert T. DeBoy, Ian T. Paulsen, Derrick E. Fouts, Jonathan A. Eisen, Sean C. Daugherty, et al. 2003. “Complete Genome Sequence of the Oral Pathogenic Bacterium Porphyromonas Gingivalis Strain W83.” Journal of Bacteriology 185 (18): 5591–5601.

Nyhus, Caitlin, Maria Pihl, Poul Hyttel, and Vanessa Jane Hall. 2019. “Evidence for Nucleolar Dysfunction in Alzheimer’s Disease.” Reviews in the Neurosciences, March. https://doi.org/10.1515/revneuro-2018-0104.

O’Leary, Nuala A., Mathew W. Wright, J. Rodney Brister, Stacy Ciufo, Diana Haddad, Rich McVeigh, Bhanu Rajput, et al. 2016. “Reference Sequence (RefSeq) Database at NCBI: Current Status, Taxonomic Expansion, and Functional Annotation.” Nucleic Acids Research 44 (D1): D733–45.

Olsen, Ingar, and Özlem Yilmaz. 2019. “Possible Role of Porphyromonas Gingivalis in Orodigestive Cancers.” Journal of Oral Microbiology 11 (1): 1563410.

Onozuka, Minoru, Kazuko Watanabe, Masafumi Fujita, Mihoko Tomida, and Satoru Ozono. 2002. “Changes in the Septohippocampal Cholinergic System Following Removal of Molar Teeth in the Aged SAMP8 Mouse.” Behavioural Brain Research 133 (2): 197–204.

Patel, Sejal, Derek Howard, Alana Man, Deborah Schwartz, Joelle Jee, Daniel Felsky, Zdenka Pausova, Tomas Paus, and Leon French. 2020. “Donor Specific Transcriptomic Analysis of Alzheimer’s Disease Associated Hypometabolism Highlights a Unique Donor, Microglia, and Ribosomal Proteins.” bioRxiv. https://doi.org/10.1101/2019.12.23.887364.

Poole, Sophie, Sim K. Singhrao, Sasanka Chukkapalli, Mercedes Rivera, Irina Velsko, Lakshmyya Kesavalu, and Stjohn Crean. 2015. “Active Invasion of Porphyromonas Gingivalis and Infection-Induced Complement Activation in ApoE-/-Mice Brains.” Journal of Alzheimer’s Disease: JAD 43 (1): 67–80.

Ray, Alpana, Deepak Kumar, Arvind Shakya, Charles R. Brown, James L. Cook, and Bimal K. Ray. 2004. “Serum Amyloid A-Activating Factor-1 (SAF-1) Transgenic Mice Are Prone to Develop a Severe Form of Inflammation-Induced Arthritis.” Journal of Immunology 173 (7): 4684–91.

Ray, Alpana, Arvind Shakya, Deepak Kumar, Merrill D. Benson, and Bimal K. Ray. 2006. “Inflammation-Responsive Transcription Factor SAF-1 Activity Is Linked to the Development of Amyloid A Amyloidosis.” Journal of Immunology 177 (4): 2601–9.

Ray, A., and B. K. Ray. 1996. “A Novel Cis-Acting Element Is Essential for Cytokine-Mediated Transcriptional Induction of the Serum Amyloid A Gene in Nonhepatic Cells.” Molecular and Cellular Biology 16 (4): 1584–94.

Roh, Jee Hoon, Hong Jiang, Mary Beth Finn, Floy R. Stewart, Thomas E. Mahan, John R. Cirrito, Ashish Heda, et al. 2014. “Potential Role of Orexin and Sleep Modulation in the Pathogenesis of Alzheimer’s Disease.” The Journal of Experimental Medicine 211 (13): 2487–96.

Sajdel-Sulkowska, E. M., and C. A. Marotta. 1984. “Alzheimer’s Disease Brain: Alterations in RNA Levels and in a Ribonuclease-Inhibitor Complex.” Science 225 (4665): 947–49.

Sawant, Kirti V., Krishna Mohan Poluri, Amit K. Dutta, Krishna Mohan Sepuru, Anna Troshkina, Roberto P. Garofalo, and Krishna Rajarathnam. 2016. “Chemokine CXCL1 Mediated Neutrophil Recruitment: Role of Glycosaminoglycan Interactions.” Scientific Reports 6 (September): 33123.

Scheff, Stephen W., Douglas A. Price, Mubeen A. Ansari, Kelly N. Roberts, Frederick A. Schmitt, Milos D. Ikonomovic, and Elliott J. Mufson. 2015. “Synaptic Change in the Posterior Cingulate Gyrus in the Progression of Alzheimer’s Disease.” Journal of Alzheimer’s Disease: JAD 43 (3): 1073–90.

Sims, N. R., D. M. Bowen, S. J. Allen, C. C. Smith, D. Neary, D. J. Thomas, and A. N. Davison. 1983. “Presynaptic Cholinergic Dysfunction in Patients with Dementia.” Journal of Neurochemistry 40 (2): 503–9.

Singhrao, Sim K., and Ingar Olsen. 2019. “Assessing the Role of Porphyromonas Gingivalis in Periodontitis to Determine a Causative Relationship with Alzheimer’s Disease.” Journal of Oral Microbiology 11 (1): 1563405.

Targońska-Stępniak, Bożena, and Maria Majdan. 2014. “Serum Amyloid A as a Marker of Persistent Inflammation and an Indicator of Cardiovascular and Renal Involvement in Patients with Rheumatoid Arthritis.” Mediators of Inflammation 2014 (November): 793628.

Trushina, Eugenia, Tumpa Dutta, Xuan-Mai T. Persson, Michelle M. Mielke, and Ronald C. Petersen. 2013. “Identification of Altered Metabolic Pathways in Plasma and CSF in Mild Cognitive Impairment and Alzheimer’s Disease Using Metabolomics.” PloS One 8 (5): e63644.

Tsujimoto, Hironori, Satoshi Ono, Takashi Majima, Nobuaki Kawarabayashi, Eiji Takayama, Manabu Kinoshita, Shuhji Seki, Hoshio Hiraide, Lyle L. Moldawer, and Hidetaka Mochizuki. 2005. “Neutrophil Elastase, MIP-2, and TLR-4 Expression during Human and Experimental Sepsis.” Shock 23 (1): 39–44.

Vemula, Pranav, Yu Jing, Hu Zhang, Jerry B. Hunt Jr, Leslie A. Sandusky-Beltran, Daniel C. Lee, and Ping Liu. 2019. “Altered Brain Arginine Metabolism in a Mouse Model of Tauopathy.” Amino Acids 51 (3): 513–28.

Vidal, R., B. Frangione, A. Rostagno, S. Mead, T. Révész, G. Plant, and J. Ghiso. 1999. “A Stop-Codon Mutation in the BRI Gene Associated with Familial British Dementia.” Nature 399 (6738): 776–81.

Vidal, R., T. Revesz, A. Rostagno, E. Kim, J. L. Holton, T. Bek, M. Bojsen-Møller, et al. 2000. “A Decamer Duplication in the 3’ Region of the BRI Gene Originates an Amyloid Peptide That Is Associated with Dementia in a Danish Kindred.” Proceedings of the National Academy of Sciences of the United States of America 97 (9): 4920–25.

Whitehouse, P. J., D. L. Price, A. W. Clark, J. T. Coyle, and M. R. DeLong. 1981. “Alzheimer Disease: Evidence for Selective Loss of Cholinergic Neurons in the Nucleus Basalis.” Annals of Neurology 10 (2): 122–26.

Whitehouse, P. J., D. L. Price, R. G. Struble, A. W. Clark, J. T. Coyle, and M. R. Delon. 1982. “Alzheimer’s Disease and Senile Dementia: Loss of Neurons in the Basal Forebrain.” Science 215 (4537): 1237–39.

Wingo, Aliza P., Wen Fan, Duc M. Duong, Ekaterina S. Gerasimov, Eric B. Dammer, Yue Liu, Nadia V. Harerimana, et al. 2020. “Shared Proteomic Effects of Cerebral Atherosclerosis and Alzheimer’s Disease on the Human Brain.” Nature Neuroscience 23 (6): 696–700.

Wu, G., and S. M. Morris Jr. 1998. “Arginine Metabolism: Nitric Oxide and beyond.” Biochemical Journal 336 (Pt 1) (November): 1–17.

Zeisel, Amit, Hannah Hochgerner, Peter Lönnerberg, Anna Johnsson, Fatima Memic, Job van der Zwan, Martin Häring, et al. 2018. “Molecular Architecture of the Mouse Nervous System.” Cell 174 (4): 999–1014.e22.

Zhang, Xiaoling, Congcong Zhu, Gary Beecham, Badri N. Vardarajan, Yiyi Ma, Daniel Lancour, John J. Farrell, et al. 2019. “A Rare Missense Variant of CASP7 Is Associated with Familial Late-Onset Alzheimer’s Disease.” Alzheimer’s & Dementia: The Journal of the Alzheimer’s Association 15 (3): 441–52.

Zhou, Qingde, Tesfahun Desta, Matthew Fenton, Dana T. Graves, and Salomon Amar. 2005. “Cytokine Profiling of Macrophages Exposed to Porphyromonas Gingivalis, Its Lipopolysaccharide, or Its FimA Protein.” Infection and Immunity 73 (2): 935–43.

